# Does sex matter in neurons’ response to hypoxic stress?

**DOI:** 10.1101/2024.10.02.616233

**Authors:** Eva J.H.F. Voogd, Marloes R. Levers, Jeannette Hofmeijer, Monica Frega, Michel J.A.M. van Putten

**Affiliations:** Department of Clinical Neurophysiology, University of Twente, Enschede, The Netherlands; Department of Neurology, Rijnstate Hospital, Arnhem, The Netherlands; Department of Neurology and Clinical Neurophysiology, Medisch Spectrum Twente, Enschede, The Netherlands

**Keywords:** Stroke, Hypoxia, Ischemic penumbra, Estrogen, Electrophysiology, Sex Characteristics, in vitro

## Abstract

**Background:** Stroke exhibits significant sex differences in incidence, response to treatment and outcome. Preclinical studies suggest that hormones, particularly estrogens, are key to differential sensitivity, as female neurons demonstrate enhanced resilience compared to males in both in vivo and in vitro models. This study investigates whether these sex-specific differences in neuronal vulnerability extend to the ischemic penumbra and explores the effects of estrogens under such conditions.

**Methods:** Primary cortical neuronal networks were generated from male and female new-born Wistar rats and cultured on micro-electrode arrays or glass coverslips. Male and female networks were subjected to hypoxic conditions, followed by a recovery phase, with or without exogenous estrogen treatment. Electrophysiological activity, including spikes and bursts, was monitored and analyzed. Apoptosis was assessed through immunocytochemistry, focusing on caspase-dependent and apoptosis inducing factor (AIF)-dependent pathways.

**Results:** Under hypoxic conditions, male and female neuronal networks exhibited a similar decrease in firing and network burst rates, with an associated increase in network burst durations. Estrogen treatment altered these dynamics, leading to increased network burst rates and decreased network burst duration for both sexes. During recovery, no significant differences were observed between estrogen-treated and untreated networks. Immunocytochemistry revealed that estrogen significantly influenced caspase-dependent apoptosis, and to a lesser extent AIF-dependent apoptosis.

**Conclusions:** In our model of the ischemic penumbra, sex-dependent differences in neuronal responses to hypoxic injury are primarily driven by estrogen, rather than intrinsic neuronal characteristics. Although our electrophysiological data demonstrated that estrogen influenced network activity, it did not offer long-term neuroprotection after hypoxia.

## 1 Introduction

Stroke is characterized by well-documented sex differences in both incidence and susceptibility to metabolic stress [1–3]. A systematic review reported that the average age at first stroke is 68.6 years for men and 72.9 years for women. Additionally, the incidence of stroke was 33% higher, and its prevalence 41% greater in men compared to women [1]. While strokes of cardio-embolic origin are more common in women, those resulting from large vessel and small vessel occlusions predominate in men [1].

Although various clinical studies report poorer outcomes of stroke in women [4], female brains may exhibit greater intrinsic resistance to hypoxic/ischemic (HI) stress compared to male brains [5]. Female neonates demonstrated reduced susceptibility to HI injury, leading to better cognitive recovery compared to males [6]. Rodent models further support these sex differences to HI injury; for instance, in vitro models using primary rat hippocampal neurons or organotypical slices, showed that male neurons exhibited a higher cell death rate than their female counterparts when exposed to hypoxia [7, 8]. Various factors are likely to contribute to this differential sensitivity to HI, including sex hormones, expression of estrogen and androgen receptors, and variations in apoptotic pathways [9].

A rich variety of preclinical studies adressed the role of sex hormones. In one of the earliest animal studies exploring the potential neuroprotective effects of estrogens, female rats underwent ovariectomy and were treated with various estrogen preparations before or after transient middle cerebral artery (MCA) occlusion. Administration of estrogen significantly reduced mortality and infarct volumes in these rats [10]. These neuroprotective effects of estrogens in experimental stroke were confirmed in other in vivo studies involving rats [11, 12]. Similar findings have been observed in other species. For example, young female mice exposed to transient MCA occlusion exhibited smaller infarct volumes than their male and postmenopausal female counterparts [13]. In addition, treatment with various steroid hormones, such as progesterone, allopregnanolone, and estradiol, reduced cell death in male and female neuronal cultures [14].

Apoptotic pathways have received significant attention, as they play a crucial role in determining cell fate following HI injury [15]. Neurons can go into apoptosis through the intrinsic, caspase-dependent, and extrinsic, Apoptosis Inducing Factor (AIF)-dependent, pathways [16]. Both in vitro and in vivo studies have shown that sensitivity to these pathways differs between sexes. Specifically, females show greater susceptibility to caspase-dependent apoptosis compared to males in response to cerebral ischemia or nitrosative stress, while in males apoptosis is more likely to be triggered by AIF-dependent pathways [17–19].

Most in vitro studies focus on acute severe ischemia, essentially simulating the infarct core. In the core, immediate severe ischemia induces rapid necrotic cell death [20]. In contrast, in regions surrounding the infarct core (i.e., the ischemic penumbra) there is some remaining perfusion from collateral arteries. Neurons in the penumbra initially remain structurally intact and potentially salvageable, making it a key target for therapies aimed at preventing irreversible brain injury [21].

In this work, we study sex specific responses to hypoxic stress in an in vitro model of the ischemic penumbra. Additionally, we assess the effects of treatment with exogenous estrogens, as well as the involvement of caspase-dependent and AIF-dependent pathways to apoptosis. To simulate conditions in the penumbral region, we use sex-segregated cultured neurons exposed to mild hypoxia. Neurons are cultured on micro-electrode arrays, for electrophysiological assessment and coverslips, for microscopical read-outs.

## 2 Methods

### 2.1 Animals

All surgical and experimental procedures regarding animal primary cell lines followed Dutch and European laws and guidelines and were approved by the Centrale Commissie Dierproeven (CCD) (AVD11000202115663). NIH principles and guidelines for reporting preclinical research were additionally used.

### 2.2 Generation of rat primary cortical neuronal networks

Neuronal networks were obtained from male and female newborn Wistar rats (P1; Janvier Labs). The sex was determined by anogenital distance [22]. Male and female rats within each litter were segregated and the brains from each sex were pooled to establish separate male and female cortical neuronal cultures. Briefly, cortices were dissociated through enzymatic treatment (0.25% trypsin) and subsequent trituration. Approximately 100,000 cells were plated on 24-well MEAs (Multi Channel Systems, Reutlingen, Germany) and approximately 80,000 cells were plated on glass coverslips. MEAs and glass coverslips were precoated with polyethylene amine (Sigma Aldrich). Cells were kept in an incubator with controlled temperature of 36 ºC, 80% humidity and 5% CO_2_ until the experiment on day in vitro (DIV) 21-25. The medium consisting of Neurobasal-glucose-pyruvate (Thermofisher, 15329741), B-27 supplement (Thermofisher, 11530536), D-glucose (1.24g/10ml), Penicillin-Streptomycin-Glutamin (100x; Thermofisher, 10378016), vitamin C (100x; Sigma Aldrich; A0278-25G), and nerve growth factor (Sigma Aldrich, N6009) was changed twice a week.

### 2.3 Experimental protocol

The experimental groups consisted of male or female neuronal networks with or without the addition of estrogen and were randomly assigned. The 24-well MEA or glass coverslips were placed in climate-controlled chambers (36 °C) that allowed for the establishment of normoxia (20% O_2_/75% N_2_/5% CO_2_) or hypoxia (2% O_2_/93% N_2_/5% CO_2_).

Stringent inclusion criteria were applied to all neuronal networks [23]. Only neuronal networks demonstrating good quality, such as sufficient cell density for proper neuron-electrode coupling and an even distribution of cells, were included.

Spontaneous neuronal network activity was recorded, which consisted of a 10-minute baseline during normoxia and then a 10-minute recording after the addition of 17*β*-estradiol. Then the hypoxia period was initiated. It took 1 hour to reach the desired oxygen level in the neuronal networks [24]. After this point, we recorded 10-minute sessions every 2 hours throughout the 36-hour hypoxia period. Following the hypoxia period, neuronal networks were returned to normoxia to assess recovery, with 10-minute recordings taken after 6 and 24 hours. To investigate the apoptotic response to hypoxia, immunocytochemical staining was applied to male and female neuronal networks at normoxia (baseline), 8, 16, 24 or 36 hours of hypoxia (Figure 1A).

**Figure 1.**
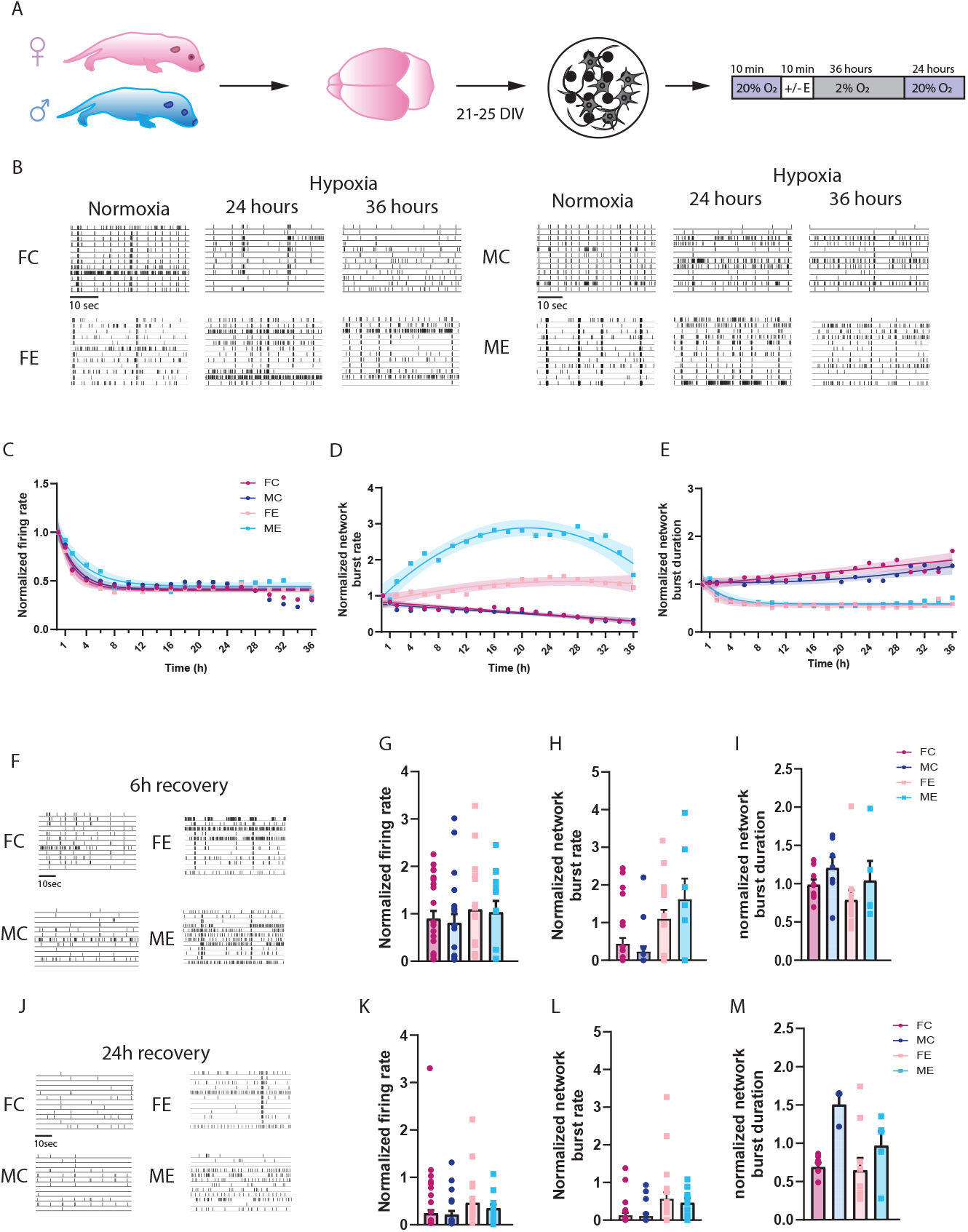
Neuronal functionality affected by hypoxia and estrogen in male and female neuronal networks. **A**. Schematic overview of generation of primary neuronal networks derived from male or female new born rat pups and experimental protocol. **B**. Representative raster plots for female control (FC), male control (MC), female estrogen (FE) and male estrogen (ME) neuronal networks during normoxia and subsequent 24 and 36 hours of hypoxia. **C-D**. Graphs showing the effect of 36 hours of hypoxia on FC (n=29), MC (n=22), FE (n=19) and ME (n=13) neuronal networks on the **C**. mean firing rate, **D**. network burst rate (NBR) and **E**. network burst duration (NBD). **F**. Representative raster plots for FC, FE, MC and ME neuronal networks during recovery where normoxia was re-established for 6 hours. **G-H**. Bar graphs showing the effect of 6 hours recovery on **G**. mean firing rate, **H**. NBR and **I**. NBD. **J**. Representative raster plots for FC, FE, MC and ME neuronal networks during recovery where normoxia was re-established for 24 hours. **K-M**. Bar graphs showing the effect of 24 hours recovery on **K**. mean firing rate, **L**. NBR and **M**. NBD. Kruskal-Wallis and Dun’s multiple comparison test was performed to test for differences between conditions shown in **G-H & K-M**.

#### 2.3.1 Exogenous estrogen treatment protocol

Neuronal networks derived from males and females were divided into estrogen-treated (E) and non-treated (C) groups. The male and female treated groups (FE and ME) received 10 nM *β*-estradiol (Merck, E2758), dissolved in 100% ethanol further diluted in culture medium, 30 minutes before the induction of hypoxia. After addition of estrogen, the experimental protocol was conducted as described above.

We initially started with a total of 120 neuronal networks on MEAs across all experimental conditions. Specifically, we used 36 male control, 36 female control, 24 male estrogen-treated, and 24 female estrogen-treated neuronal networks. During the experimental process, some neuronal networks were excluded based on the pre-established inclusion criteria. As a result, the final number of neuronal networks on MEAs included in the analysis was 22 for MC, 29 for FC, 13 for ME, and 19 for FE.

We initially started with a total of 120 neuronal networks on coverslips across all experimental conditions. Specifically, we used 32 male control, with 8 networks assigned to each time point (normoxia, 8h, 16h, 24h and 36h of hypoxia), and similarly 32 female control. We used 16 neuronal networks for both male estrogen-treated and female estrogen-treated, with 4 networks assigned to each time point. During the experimental process, some neuronal networks were excluded based on the pre-established inclusion criteria. As a result, the final number of neuronal networks on coverslips are described in supplementary table 8.

### 2.4 Electrophysiology

Electrophysiological activity was assessed utilizing the Multiwell-MEA system (Multi Channel Systems, Reutlingen, Germany) and recorded with the Multiwell-screen software. Signals were sampled at 10 kHz after high-pass (2nd order Butterworth filter, 100 Hz) and low-pass (4th order Butterworth filter, 3.5 kHz) filtering.

Data analysis was performed using a combination of Multiwell Analyzer software (Multi Channel Systems, Reutlingen, Germany) and custom MATLAB scripts (The MathWorks, Natick, MA, USA).

#### 2.4.1 Spike detection and firing rate

Spike events were identified when the signal exceeded baseline noise by 5 times the standard deviation. An electrode was deemed active if it displayed a minimum of 0.1 spikes per second. Subsequently, the mean firing rate was computed by determining the spike count per electrode over time and averaging across all active electrodes per well.

#### 2.4.2 Burst detection

Single-channel bursts were identified as sequences of spikes containing a minimum of 4 spikes, each with an inter-spike interval of 50 ms or less. A minimum interval of 100 ms was used to differentiate between consecutive bursts. A channel was categorized as a bursting channel if it displayed a frequency of at least 0.4 bursts per minute.

#### 2.4.3 Quantification of network bursts

A network burst was defined as a sequence of temporally overlapping bursts from at least six distinct channels, with at least four channels showing overlapping bursting activity during the sequence. Following detection, network bursts were characterized by burst rate and average burst duration.

For neuronal networks to be deemed active and included in the analysis, neuronal networks needed to display a mean firing rate (MFR) greater than *>* 0.1 spikes per second and at least 1 network burst per minute.

### 2.5 Immunocytochemistry

#### 2.5.1 Caspase-dependent apoptosis

Caspase-dependent apoptosis was assessed through immunocytochemical staining for caspase 3/7. Cell Event (1:500, Thermoscientific) was added to the cultures and cultures were incubated for 30 min at 37 °C to stain the cells expressing caspase 3/7. Next, the cells were subjected to different experimental conditions. At the end of the experiments, cells were carefully washed with Dulbecco’s phosphate-buffered saline (dPBS, Gibco) and fixated with 3.7% paraformaldehyde (PFA, Sigma Aldrich) for 15 min at room temperature (RT). Lastly, nuclei were stained with DAPI (1:7500, Sigma Aldrich) for 10 min at RT. The samples were carefully washed again, mounted with Mowiol, and stored at 4 °C.

#### 2.5.2 Apoptosis Inducing Factor-dependent apoptosis

AIF-dependent apoptosis was assessed by immunocytochemical staining for AIF. Primary cortical neuronal networks that had been subjected to different experimental conditions, were fixed using 3.7% PFA for 15 minutes at RT, then rinsed with dPBS and stored in dPBS at 4°C until staining. To begin, samples were permeabilized using 0.2% triton X-100 (Sigma Aldrich) in dPBS and washed with dPBS. Subsequently, a blocking buffer (2% bovine serum albumin, Sigma-Aldrich, in dPBS) was applied for 30 minutes in RT to inhibit nonspecific binding. Cultures were stained with mouse anti-AIF (1:1000, M4403-50, Sigma Aldrich) diluted in blocking buffer and incubated overnight at 4 °C. After thorough washing with dPBS, a secondary antibody, goat anti-mouse (AF647, 1:2000, A11029, Invitrogen), was applied for 1 hour at RT. Subsequently, the nuclei were stained with DAPI (1:7500) for 10 minutes at RT. After another round of careful washing, the samples were mounted with Mowiol, dried, and stored at 4 °C.

### 2.6 Microscopy

Images of ten random fields per neuronal network were captured at 40x magnification using a Nikon Eclipse 50i Epi-Fluorescence microscope (Nikon, Japan).

To investigate the caspase-dependent pathway, image analysis was conducted using custom MATLAB scripts to preprocess the captured images. Subsequently, a blind manual cell count was performed, distinguishing cells positive for caspase 3/7 (green fluorescence) from cells positive for DAPI (staining all cell nuclei). The ratio of apoptotic cells was calculated by dividing the count of caspase-positive cells by the total count of DAPI-positive cells. This rigorous approach ensured an unbiased evaluation of caspase-dependent pathway activation in response to hypoxic stress within neuronal networks.

To investigate the AIF-dependent pathway, custom MATLAB scripts were used to preprocess the images and compute the mean intensity of AIF in each image. The mean intensity was corrected for the total amount of DAPI positive cells per image.

### 2.7 Statistics

To model the temporal evolution of our electrophysiological and immunocytochemical readouts, we used first and second order polynomials and a single exponential function. These models effectively capture linear, curvilinear, and exponential trends, as commonly observed in natural systems [25,26], including neuronal survival and electrophysiological responses during hypoxic stress. Model selection was based on the Akaike information criterium (AIC) [27]. A difference of the AIC larger than two was used to consider models significantly different [28, 29]. If model selection for different experimental conditions resulted in the same model, we subsequently tested for significant differences in the parameter values. If data were normally distributed (assessed using the Shapiro-Wilk test) t-statistics were used to compare parameters, otherwise the Mann-Whitney U test.

Additionally, we performed an analysis of neuronal recovery, at 6 and 24 hours after exposure for 36 hours to low oxygen conditions, using Kruskal-Wallis and Dun’s multiple comparison tests.

Identification of outliers was performed employing Robust Regression and Outlier removal (ROUT) method with Q = 1% leading to the removal of data points in the lower and upper 1% of the normal distribution. Data are presented as mean *±* standard error of the mean (SEM). Statistical analyses were conducted using Graphpad Prism 10 (GraphPad Software, 15 Inc., California, USA).

## 3 Results

### 3.1 Electrophysiology

After three weeks in vitro, primary cortical neurons derived from male and female rat pups exhibited rich spontaneous dynamical behavior characterized by various patterns of electrical activity, consisting of spikes, bursts, and network bursts (Fig. 1B).

#### 3.1.1 Male vs. female neuronal networks

Under hypoxic conditions, the firing rates of male and female control neuronal networks decreased and reached a plateau. The neuronal networks of both sexes followed first-order exponential decay kinetics (see supp. table 1 for AIC criteria), with time constants of the order of 2.5 h, eventually reaching a mean value of approximately 45% of baseline firing rate (Fig. 1C). Concurrently, the network burst rate (NBR) decreased while the network burst duration (NBD) increased, for both male and female networks. NBR of both sexes was best described with a linear model (see supp. table 2 for AIC criteria). The slopes of NBR models between male and female neuronal networks were not significantly different (p=0.9). However, for male networks, NBD was best described by a quadratic function, while for female networks, NBD was best described by a linear function (supp. table 4 & 5).

#### 3.1.2 Effect of estrogen on male and female neuronal networks

The network firing rates of male and female neurons exposed to estrogen decreased in a similar way as controls (p=0.11), showing an exponential decrease with a comparable time constant (Fig 1C). In contrast, the NBRs of male and female networks treated with estrogen increased, with their evolution best described by a quadratic function (see supp. table 2 for model choice based on AIC). NBDs, on the other hand, decreased and reached a plateau. These were best modeled as exponential behavior (Fig. 1E). These differences in models reflect statistically significant differences in dynamics between estrogen-treated neurons and control neurons. Furthermore, the NBR in male networks treated with estrogen increased significantly more compared to female networks treated with estrogen. The NBD was not significantly different in male and female neurons treated with estrogen (supp. table 3).

Following the hypoxic period, normoxia was restored and the recovery phase commenced. After 6 hours of recovery, firing rates and NBDs returned close to baseline values, while NBR did not. After 24 hours, the firing rate of controls decreased, while estrogen-treated networks stayed relatively stable. At 24 hours the NBR was similar compared to 6 hours recovery. We found no significant differences in the firing rates, NBR and NBD between all four conditions at 6 or 24 hours after recovery (Fig. 1F-M).

### 3.2 Immunocytochemistry

#### 3.2.1 Male vs. female neuronal networks

Under hypoxic conditions, caspase 3/7 activity in male and female control neuronal networks increased over time, following a linear temporal evolution (See supp. table 6 for model choice based on AIC; Fig. 2A-C). No significant differences were found in slopes between male and female neuronal networks (p=0.37).

**Figure 2.**
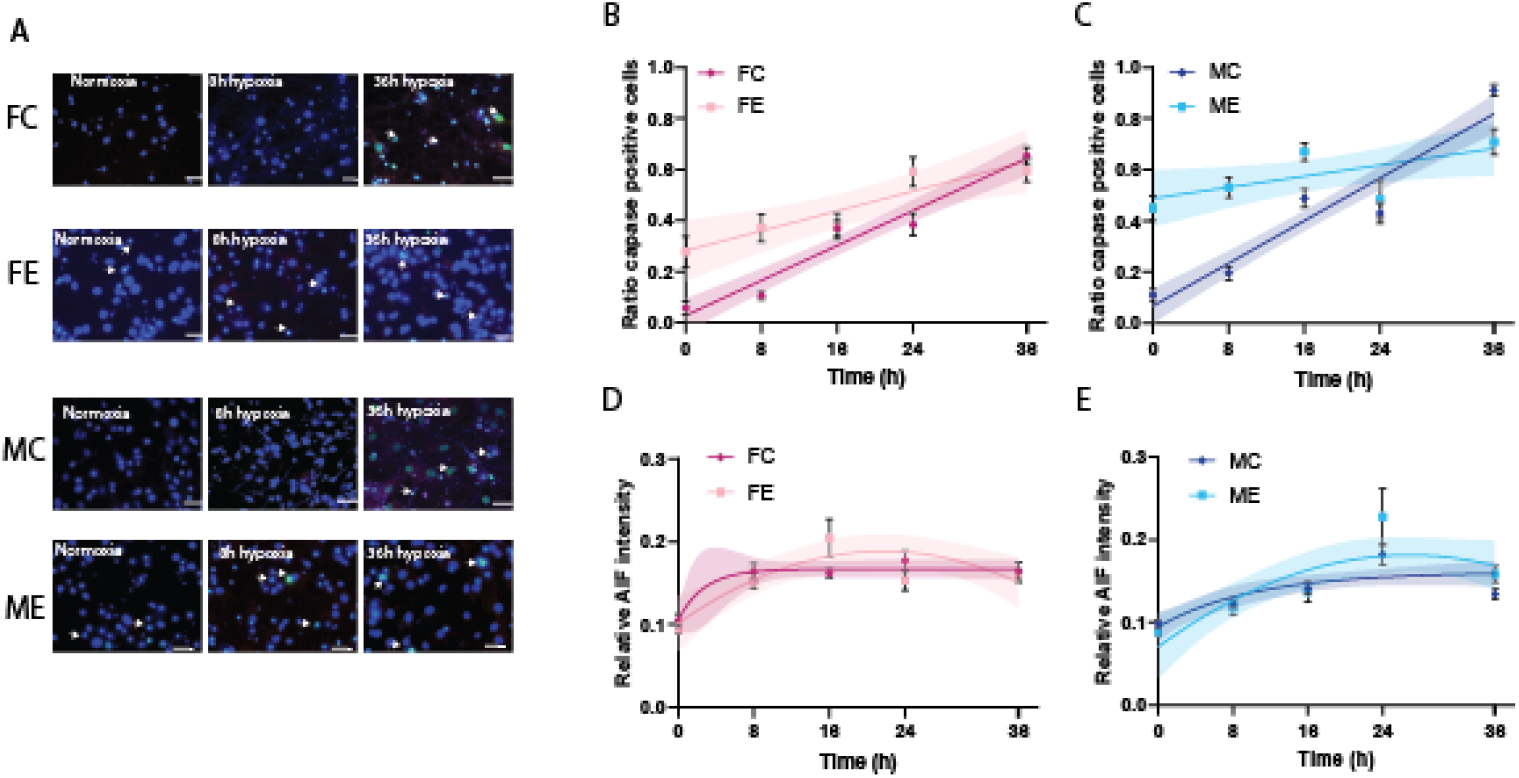
Different modes of apoptosis in male and female neuronal networks under hypoxic stress. **A**. Representative images of female control (FC; n=5-8), female estrogen-treated (FE; n=2-4), male control (MC; n=6-8) and male estrogen-treated (ME; n=2-4) neuronal networks showing caspase-dependent (bright blue/green, arrows) and AIF-dependent (red) apoptosis with nuclei positive for DAPI (blue) in normoxia, 8 and 36 hours of hypoxia. Arrows indicate caspase-positive cells (scale bar = 30 *μ*m). **B, C**. Linear regression analyses of the ratio of caspase-positive cells of **B**. female control (FC; dark pink) and female estrogen-treated (FE; light pink) and **C**. male control (MC; dark blue) and male estrogentreated (ME; light blue). **D, E**. graphs of curves of relative AIF intensity for **D**. female control (FC; dark pink; exponential) and female estrogen-treated (FE; light pink; quadratic) and **E**. male control (MC; dark blue; exponential) and male estrogen-treated (ME; light blue; quadratic).

Under hypoxic conditions, AIF activity in control conditions increased and reached a plateau, exhibiting exponential dynamics (Fig. 2D & E). The curves of female and male control neurons were significantly different (p=0.005).

#### 3.2.2 Effect of estrogen on male and female neuronal networks

After estrogen treatment, baseline values of caspase 3/7 activity were increased, but the slopes of the temporal evolution of caspase 3/7 activity were significantly reduced compared to controls in males (p*<*0.0001) and in females (p=0.0013; Fig. 2B-C).

AIF activity in estrogen-treated groups initially increased followed by a decrease and was best described by a quadratic function (see supp. table 7 for model choice based on AIC; Fig. 2D & E). These differences in models reflect statistically significant differences in dynamics between estrogen-treated neurons and control conditions.

## 4 Discussion

We studied the response of male and female cultured neuronal networks to mild transient hypoxia. Our findings reveal no intrinsic differences in electrophysiological response or caspase-dependent apoptosis between male and female neurons. However, estrogen clearly modulated the neuronal responses to hypoxic stress in both sexes, increasing network burst rates. Despite this modulation during hypoxia, we found not evidence for long-term neuroprotection by estrogen.

### Electrophysiology

The gradual decrease in mean firing rate to a new plateau under hypoxic conditions reflects compromised neuronal function in the ischemic penumbra where sufficient energy remains to sustain some residual neuronal activity for several hours. This was accompanied by changes in synchronization, evidenced by a decrease in network burst rate and an increase in the average network burst duration. These changes primarily reflect selective synaptic failure, a critical process in the penumbral region, where neuronal function is compromised without significant depolarization or swelling [30, 31]. Reduced spiking due to impaired synaptic transmission has also been observed in vivo in rats subjected to MCA occlusion [32]. None of our three electrophysiological readouts during hypoxia showed sex differences, suggesting that intrinsic neuronal responses to hypoxic stress are similar in both males and females.

In contrast, in neuronal networks treated with exogenous 17*β*-estradiol during hypoxia, synchronization as reflected by the network burst rate and duration, was significantly different from untreated neuronal networks, with an increase in network burst rate and a decrease in network burst duration. The changes in network burst rate were more pronounced in males than in females. A similar sex-dependent effect was observed in a study in rats, where estrogen administration increased the CA1 population spike amplitude in male slices under normoxia but slightly depressed it in females [33]. These differences possibly result from different expression of estrogen receptors [34]. Previous research has shown that 17*β*-estradiol influences calcium homeostasis and modulates synaptic transmission, potentially preserving network behavior [35–37]. Synaptic transmission can be enhanced by the effect of estrogen on the amplitude and frequency of mini postsynaptic currents and the generation of new synapses [35]. Additionally, 17*β*-estradiol potentiates synaptic transmission through AMPA and NMDA receptors, while attenuating GABA-mediated inhibition under normoxic conditions [36, 37]. Our observed effects of estrogen on network synchronization during hypoxia may result from these specific effects on excitatory and inhibitory synapses.

Despite the pronounced effect of estrogen on the temporal evolution of the network burst rate and duration during hypoxia, these effects were not sustained during the recovery phase. After 6 and 24 hours of recovery, no significant differences in any of the readouts remained, including network firing rates. This suggests that estrogen did not provide long-term neuroprotection in our model of the ischemic penumbra. This contrasts with previous studies showing improved cell viability with estrogen in vitro [38, 39]. Differences in extracellular matrix (ECM) composition between in vitro and in vivo models may partly explain the differences in effects of estrogen. In cultured neurons, the ECM is more extensive than in the intact brain, which could influence how neurons respond to hypoxic stress and treatment. In the brain, cell swelling and subsequent cell death are probably key drivers of the transition towards irreversible injury in core and penumbra [40], while isolated synaptic failure also plays a role, especially in the penumbra. Under in vitro conditions the timelines or mechanisms towards neuronal death are probably different. Moreover, the method of inducing metabolic stress differed between studies, with nitrosative stress, oxygen-glucose deprivation, or excitotoxicity and differences in depth and duration [38, 41]. These variations may contribute to the observed differences in effects of estrogen across models and studies.

### Immunocytochemistry

Caspase and AIF are key markers of apoptosis, the primary pathway towards cell death in the ischemic penumbra [42, 43]. Under control conditions, caspase 3/7 activity increased linearly over time in both male and female networks, indicating progressive apoptosis. In estrogen-treated networks, caspase 3/7 activity was elevated in both sexes before induction of hypoxia, possibly reflecting estrogen’s modulation of this apoptotic pathway [44, 45]. Estrogen, particularly through estrogen receptor *β*, influences apoptotic signaling in neurons [46].

Differences in AIF activity between male and female neuronal networks, although significant, were relatively small, with a variation of less than 10-15%. Sex-specific AIF activation has been reported under metabolic stress, with male neurons showing more pronounced AIF release and nuclear translocation compared to females [17, 47, 48]. However, our study did not find convincing differences in estrogen-mediated modulation of AIF activity during hypoxia, which may be due to the milder transient hypoxia used, compared to the more severe stress models such as oxygen-glucose deprivation, nitrosative stress, and excitotoxicity used in previous studies [17, 47, 48].

Given that no differences in neuronal functionality persisted after recovery between all conditions, we did not further investigate apoptotic factors post-hypoxia.

### Linking electrophysiology and immunocytochemistry

Our electrophysiological findings highlight functional changes in neuronal networks under hypoxia, both with and without estrogen. In addition, our immunocytochemical results offer insights into the underlying apoptotic processes. We observed increased caspase 3/7 activity under normoxia with estrogen, while in hypoxia, estrogen led to changes in network synchronization. Synaptic activity plays a role in promoting resistance to apoptosis [49, 50]. Estrogen’s enhancement of synaptic transmission may help maintain neuronal network function and inhibit apoptotic pathway activation under hypoxic stress. We propose that estrogen’s effects on neuronal functionality may be mediated indirectly through the preservation of synaptic integrity and function, rather than through direct action on cell survival pathways. Supporting this, synaptic activity has been shown to inhibit the caspase-dependent apoptotic pathway, reducing apoptosis and promoting cell survival [49].

### Limitations

Our experimental design focused exclusively on hypoxia-induced neuronal responses and did not include other forms of metabolic stress, such as glucose deprivation. While hypoxia is a critical component of ischemic stroke pathology, the exclusion of additional stressors may limit the comprehensiveness of our conclusions. Secondly, the choice of an animal model system presents another potential limitation. We utilized primary cortical neurons derived from rats, rather than human-derived neuronal networks. Although rat neurons provide valuable information on basic cellular processes and physiological responses, there may be inherent differences in neuronal function and susceptibility to hypoxic injury between rodents and humans [40]. Finally, we considered a limited number of models to describe the temporal changes in our observations. Although the fits were satisfactory, the models or the differences between them do not provide detailed information about the underlying biology.

## Conclusion

In our model of the ischemic penumbra, we did not observe sex-dependent responses to metabolic stress under control conditions. Estrogen differentially affected the synchrony of male and female neuronal networks, but no differences in neuronal functionality were observed after restoration of normoxia. Our results suggest that reported differences in neuronal vulnerability between sexes most likely result from estrogen rather than intrinsic neuronal properties.

## References

[1] P. Appelros, B. Stegmayr, and A. Terent. Sex differences in stroke epidemiology: a systematic review. Stroke, 40(4):1082–90, 2009.

[2] D. Mozaffarian, E.J. Benjamin, and et al. Heart disease and stroke statistics–2015 update: a report from the american heart association. Circulation, 131(4):e29–322, 2015.

[3] M. J. Reeves, G. Bushnell, C. D.and Howard, J. W. Gargano, P. W. Duncan, Khatiwoda A. Lynch, G., and L. Lisabeth. Sex differences in stroke: epidemiology, clinical presentation, medical care, and outcomes. Lancet Neurology, 7(10):915–926, 2008.

[4] M. Xu, A. Amarilla Vallejo, C. Cantalapiedra Calvete, A. Rudd, C. Wolfe, M.D.L. O’Connell, and A. Douiri. Stroke outcomes in women: A population-based cohort study. Stroke, 53(10):3072–3081, 2022.

[5] V. Chanana, M. Hackett, N. Deveci, N. Aycan, B. Ozaydin, N.S. Cagatay, D. Hanalioglu, D.B. Kintner, K. Corcoran, S. Yapici, F. Camci, J. Eickhoff, K.M. Frick, P. Ferrazzano, J.E. Levine, and P. Cengiz. Trkb-mediated sustained neuroprotection is sex-specific and erα-dependent in adult mice following neonatal hypoxia ischemia. Biology of Sex Differences, 15(1):1, 2024.

[6] M.D. Lauterbach, S. Raz, and C.J. Sander. Neonatal hypoxic risk in preterm birth infants: the influence of sex and severity of respiratory distress on cognitive recovery. Neuropsychology, 15(3):411–420, 2001.

[7] H. Li, S. Pin, Z. Zeng, M. M. Wang, K. A. Andreasson, and L. D. McCullough. Sex differences in cell death. Ann Neurol, 58(2):317–21, 2005.

[8] A. Heyer, M. Hasselblatt, N. von Ahsen, H. Hafner, A. L. Siren, and H. Ehrenreich. In vitro gender differences in neuronal survival on hypoxia and in 17beta-estradiol-mediated neuroprotection. J Cereb Blood Flow Metab, 25(4):427–30, 2005.

[9] S.L. Zup, N.S. Edwards, and M.M. McCarthy. Sex- and age-dependent effects of androgens on glutamate-induced cell death and intracellular calcium regulation in the developing hippocampus. Neuroscience, 281:77–87, 2014.

[10] J. W. Simpkins, G. Rajakumar, Y. Q. Zhang, C. E. Simpkins, D. Greenwald, C. J. Yu, N. Bodor, and A. L. Day. Estrogens may reduce mortality and ischemic damage caused by middle cerebral artery occlusion in the female rat. Journal of neurosurgery, 87(5):724–30, 1997.

[11] D. B. Dubal, M. L. Kashon, L. C. Pettigrew, J. M. Ren, S. P. Finklestein, S. W. Rau, and P. M. Wise. Estradiol protects against ischemic injury. Journal of cerebral blood flow and metabolism, 18(11):1253–8, 1998.

[12] Y.Q. Zhang, J. Shi, G. Rajakumar, A.L. Day, and J.W. Simpkins. Effects of gender and estradiol treatment on focal brain ischemia. Brain Research, 784(1):321–324, 1998.

[13] B. Manwani, F. Liu, V. Scranton, M. D. Hammond, L. H. Sansing, and L. D. McCullough. Differential effects of aging and sex on stroke induced inflammation across the lifespan. Exp Neurol, 249:120–31, 2013.

[14] R. Altaee and C. L. Gibson. Sexual dimorphism following in vitro ischemia in the response to neurosteroids and mechanisms of injury. BMC Neuroscience, 21(1):5, January 2020.

[15] J. T. Lang and L. D. McCullough. Pathways to ischemic neuronal cell death: are sex differences relevant? Journal of Translational Medicine, 6(1):33, June 2008.

[16] S. K. Rupinder, A. K. Gurpreet, and S. Manjeet. Cell suicide and caspases. Vascul Pharmacol, 46(6):383–93, 2007.

[17] L. Du, H. Bayir, Y. Lai, X. Zhang, P. M. Kochanek, S. C. Watkins, S. H. Graham, and R. S. Clark. Innate gender-based proclivity in response to cytotoxicity and programmed cell death pathway. J Biol Chem, 279(37):38563–70, 2004.

[18] L. D. McCullough, Z. Zeng, K. K. Blizzard, I. Debchoudhury, and P. D. Hurn. Ischemic nitric oxide and poly (adp-ribose) polymerase-1 in cerebral ischemia: male toxicity, female protection. J Cereb Blood Flow Metab, 25(4):502–12, 2005.

[19] S. Renolleau, S. Fau, C. Goyenvalle, L. M. Joly, D. Chauvier, E. Jacotot, J. Mariani, and C. Charriaut-Marlangue. Specific caspase inhibitor q-vd-oph prevents neonatal stroke in p7 rat: a role for gender. J Neurochem, 100(4):1062–71, 2007.

[20] O. Mărgăritescu, L. Mogoantă, I. Pirici, D. Pirici, D. Cernea, and C. Mărgăritescu. Histopathological changes in acute ischemic stroke. Romanian Journal of Morphology and Embryology, 50(3):327–339, 2009.

[21] J. Hofmeijer and M. J. A. M. van Putten. Ischemic cerebral damage: an appraisal of synaptic failure. Stroke, 43(2):607–615, 2012. Epub 2011 Dec 29.

[22] M. S. Marty, R. E. Chapin, L. G. Parks, and B. A. Thorsrud. Development and maturation of the male reproductive system. Birth Defects Research Part B: Developmental and Reproductive Toxicology, 68(2):125–136, April 2003.

[23] B. Mossink, A.H.A Verboven, E.J.H. van Hugte, T. M. Klein Gunnewiek, G. Parodi, K. Linda, C. Schoenmaker, T. Kleefstra, T. Kozicz, H. Bokhoven, D. Schubert, N. Nadif Kasri, and M. Frega. Human neuronal networks on micro-electrode arrays are a highly robust tool to study disease-specific genotype-phenotype correlations in vitro. Stem Cell Reports, 16(9):2182–2196, 2021.

[24] S. Pires Monteiro, E. Voogd, L. Muzzi, G. De Vecchis, B. Mossink, M. Levers, G. Hassink, M. Van Putten, J. Le Feber, J. Hofmeijer, and M. Frega. Neuroprotective effect of hypoxic preconditioning and neuronal activation in a in vitro human model of the ischemic penumbra. Journal of Neural Engineering, 18(3):036016, 2021.

[25] R. E. Ricklefs and G. L. Miller. Ecology. W. H. Freeman and Company, 2000.

[26] Oliver Schabenberger and Francis J. Pierce. Contemporary Statistical Models for the Plant and Soil Sciences. CRC Press, Boca Raton, 1st edition edition, 2001.

[27] A. Tsoularis and J. Wallace. Analysis of logistic growth models. Mathematical Biosciences, 179(1):21–55, 2002.

[28] H. Akaike. A new look at the statistical model identification. IEEE Transactions on Automatic Control, 19(6):716–723, December 1974.

[29] K. P. Burnham and D. R. Anderson. Basic use of the information–theoretic approach. In Model Selection and Multimodel Inference, pages 98–148. Springer, New York, 2004.

[30] S. Fedorovich, J. Hofmeijer, M.J. van Putten, and J. le Feber. Reduced synaptic vesicle recycling during hypoxia in cultured cortical neurons. Frontiers in Cellular Neuroscience, 11:32, Feb 2017.

[31] I. Zironi and G. Aicardi. Hypoxia depresses synaptic transmission in the primary motor cortex of the infant rat—role of adenosine a1 receptors and nitric oxide. Biomedicines, 10:2875, 2022.

[32] H. Bolay, Y. Gürsoy-Ozdemir, Y. Sara, R. Onur, A. Can, and T. Dalkara. Persistent defect in transmitter release and synapsin phosphorylation in cerebral cortex after transient moderate ischemic injury. Stroke, 33:1369–1375, 2002.

[33] T. Teyler, R. Vardaris, D. Lewis, and A. Rawitch. Gonadal steroids: effects on excitability of hippocampal pyramidal cells. Science, 209(4460):1017–1018, 1980.

[34] N.H. Contoreggi, S. Mazid, L.B. Goldstein, J. Park, A.C. Ovalles, E.M. Waters, M.J. Glass, and T.A. Milner. Sex and age influence gonadal steroid hormone receptor distributions relative to estrogen receptor β-containing neurons in the mouse hypothalamic paraventricular nucleus. The Journal of Comparative Neurology, 529(7):1522–1543, Apr 2021.

[35] R. Hu, W. Cai, X. Wu, and Z. Yang. Astrocyte-derived estrogen enhances synapse formation and synaptic transmission between cultured neonatal rat cortical neurons. Neuroscience, 144:1229–1240, 2007.

[36] C. S. Woolley. Acute effects of estrogen on neuronal physiology. Annual Review of Pharmacology and Toxicology, 47:657–680, 2007.

[37] M. T. Kim, W. Soussou, G. Gholmieh, A. Ahuja, A. Tanguay, T. W. Berger, and R. D. Brinton. 17beta-estradiol potentiates field excitatory postsynaptic potentials within each subfield of the hippocampus with greatest potentiation of the associational/commissural afferents of ca3. Neuroscience, 141(1):391–406, 2006. Epub 2006 May 24.

[38] C. Harms, M. Lautenschlager, A. Bergk, J. Katchanov, D. Freyer, K. Kapinya, U. Herwig, D. Megow, U. Dirnagl, J. R. Weber, and H. Hörtnagl. Differential mechanisms of neuroprotection by 17 beta-estradiol in apoptotic versus necrotic neurodegeneration. Journal of Neuroscience, 21(8):2600–2609, 2001.

[39] J. Morán, M. Perez-Basterrechea, P. Garrido, et al. Effects of estrogen and phytoestrogen treatment on an in vitro model of recurrent stroke on ht22 neuronal cell line. Cellular and Molecular Neurobiology, 37:405–416, 2017.

[40] B.D. Semple, K. Blomgren, K. Gimlin, D.M. Ferriero, and L.J. Noble-Haeusslein. Brain development in rodents and humans: Identifying benchmarks of maturation and vulnerability to injury across species. Progress in Neurobiology, 106-107:1–16, Jul-Aug 2013.

[41] P. Duong, M.A.A. Tenkorang, J. Trieu, C. McCuiston, N. Rybalchenko, and R.L. Cunningham. Neuroprotective and neurotoxic outcomes of androgens and estrogens in an oxidative stress environment. Biology of Sex Differences, 11(1):12, 2020.

[42] M.D. Linnik, R.H. Zobrist, and M.D. Hatfield. Evidence supporting a role for programmed cell death in focal cerebral ischemia in rats. Stroke, 24(12):2002–2008, 1993.

[43] C. Charriaut-Marlangue, I. Margaill, F. Borrega, M. Plotkine, and Y. Ben-Ari. Ngnitro-l-arginine methyl ester reduces necrotic but not apoptotic cell death induced by reversible focal ischemia in rat. European Journal of Pharmacology, 310(2-3):137–140, 1996.

[44] J. Nilsen, G. Mor, and F. Naftolin. Estrogen-regulated developmental neuronal apoptosis is determined by estrogen receptor subtype and the fas/fas ligand system. Journal of Neurobiology, 43(1):64–78, 2000.

[45] D. B. Dubal, S. W. Rau, P. J. Shughrue, H. Zhu, J. Yu, A. B. Cashion, S. Suzuki, L. M. Gerhold, M. B. Bottner, I. Merchanthaler, M. S. Kindy, and P. M. Wise. Differential modulation of estrogen receptors (ers) in ischemic brain injury: a role for eralpha in estradiol-mediated protection against delayed cell death. Endocrinology, 147(6):3076–3084, 2006.

[46] L. Zhao and R.D. Brinton. Estrogen receptor alpha and beta differentially regulate intracellular ca(2+) dynamics leading to erk phosphorylation and estrogen neuroprotection in hippocampal neurons. Brain Research, 1172:48–59, 2007.

[47] J. Sharma, G. Nelluru, M. A. Wilson, M. V. Johnston, and M. A. Hossain. Sex-specific activation of cell death signalling pathways in cerebellar granule neurons exposed to oxygen glucose deprivation followed by reoxygenation. ASN Neuro, 3(2), 2011.

[48] H. Y. Cheng, S. H. Hung, and P. J. Chu. Rescue from sexually dimorphic neuronal cell death by estradiol and pi3 kinase activity. Cellular and Molecular Neurobiology, 36:767–775, 2016.

[49] F. Leveille, S. Papadia, M. Fricker, K. F. S. Bell, F. X. Soriano, M.-A. Martel, and G. E. Hardingham. Suppression of the intrinsic apoptosis pathway by synaptic activity. Journal of Neuroscience, 30(7):2623–2635, 2010.

[50] M. Catsicas, Y. Péquignot, and P.G. Clarke. Rapid onset of neuronal death induced by blockade of either axoplasmic transport or action potentials in afferent fibers during brain development. Journal of Neuroscience, 12(12):4642–4650, 1992.

