## Supplementary materials 1 for "Does sex matter in neurons’ response to hypoxic stress?"

### 5 Supplementary Material

- **One-Phase Exponential Decay:** We fitted our data to a one-phase exponential decay model to characterize the rate of decrease over time. This model assumes the form:

$$f(x) = (a_0 - b) * exp(-t/\tau) + b \quad (1)$$

  where $a_0$ is the initial value, $\tau$ is the decay constant, and $b$ is the asymptotic value. This approach was used to describe how the data approached a stable plateau following an initial rapid change.

- **Linear Regression:** For datasets exhibiting a linear trend, we applied linear regression. The model used was:

$$f(x) = a + bx \quad (2)$$

  where $\beta_0$ represents the intercept and $\beta_1$ the slope. This model was utilized to analyze linear relationships between time and the dependent variable.

- **Second-Order Polynomial Regression:** To capture non-linear trends, we fitted a second-order polynomial regression model of the form:

$$f(x) = ax^2 + bx + c \quad (3)$$

  where $\alpha_0$, $\alpha_1$, and $\alpha_2$ are the coefficients of the polynomial. This model was used to explore quadratic relationships and identify patterns not captured by linear models.

Table 1: Overview of Best Models, AIC Values, and Parameter Estimates for Each Condition, mean firing rate

| **Condition** | **Best Model** | **AIC** | **Parameter Estimates** |
|---|---|---|---|
| MC | Exponential | -120.351 | [0.603, 2.634, 0.402] |
| FC | Exponential | -97.949 | [0.606, 2.389, 0.401] |
| ME | Exponential | -141.302 | [0.583, 3.654, 0.445] |
| FE | Exponential | -118.194 | [0.613, 1.786, 0.405] |

Table 2: Overview of Best Models, AIC Values, and Parameter Estimates for Each Condition, network burst rate

| **Condition** | **Best Model** | **AIC** | **Parameter Estimates** |
|---|---|---|---|
| MC | Linear | -115.233 | [-0.015, 0.833] |
| FC | Linear | -101.603 | [-0.016, 0.882] |
| ME | Quadratic | -68.250 | [-0.004, 0.177, 1.031] |
| FE | Quadratic | -86.503 | [-0.001, 0.033, 0.927] |

Table 3: Comparison of Parameters Between ME and FE Conditions, network burst rate

| Parameter | ME | FE | t-statistic | p-value |
|---|---|---|---|---|
| a3 | -1.719 | -0.560 | -4.067 | 0.001 |
| b3 | 4.728 | 18.491 | -0.944 | 0.357 |
| c3 | 2.597 | 1.511 | 4.835 | 0.0001 |

Table 4: Overview of Best Models, AIC Values, and Parameter Estimates for Each Condition, network burst duration

| Condition | Best Model | AIC | Parameter Estimates |
|---|---|---|---|
| MC | Quadratic | -110.272 | [0.0, -0.007, 1.074] |
| FC | Linear | -92.914 | [0.012, 1.021] |
| ME | Exponential | -105.263 | [0.567, 2.723, 0.586] |
| FE | Exponential | -113.797 | [0.518, 2.309, 0.551] |

Table 5: Comparison of Parameters Between MC and FC Condition, network burst duration

| Parameter | MC | FC | t-statistic | p-value |
|---|---|---|---|---|
| a3 | 0.567 | 0.518 | 0.614 | 0.546 |
| b3 | 2.723 | 2.309 | 0.430 | 0.672 |
| c3 | 0.586 | 0.551 | 1.292 | 0.212 |

Table 6: Overview of Best Models, AIC Values, and Parameter Estimates for Each Condition, Caspase activity

| Condition | Best Model | AIC | Parameter Estimates |
|---|---|---|---|
| MC | Linear | -26.679 | [0.01, 0.283] |
| FC | Linear | -26.261 | [0.017, 0.031] |
| ME | Linear | -21.765 | [0.006, 0.475] |
| FE | Linear | -20.426 | [0.021, 0.07] |

Table 7: Overview of Best Models, AIC Values, and Parameter Estimates for Each Condition, AIF activity

| Condition | Best Model | AIC | Parameter Estimates |
|---|---|---|---|
| MC | Exponential | -33.825 | [-0.076, 3.851, 0.17] |
| FC | Exponential | -46.868 | [-0.063, 3.132, 0.168] |
| ME | Quadratic | -30.442 | [-0.0, 0.008, 0.076] |
| FE | Quadratic | -38.056 | [-0.0, 0.006, 0.091] |

Table 8: Number of included neuronal networks for immunocytochemical analyses.

| Condition | 0h | 8h | 16h | 24h | 36h |
|---|---|---|---|---|---|
| MC (n=) | 6 | 8 | 7 | 8 | 6 |
| FC (n=) | 5 | 7 | 8 | 6 | 7 |
| ME (n=) | 2 | 4 | 4 | 3 | 4 |
| FE (n=) | 2 | 4 | 4 | 4 | 3 |

### Core Set of Reporting Standards for Rigorous Study Design

#### Randomization

- Animals should be assigned randomly to the various experimental groups, and the method of randomization reported. **The experimental groups consisted of male or female neuronal networks with or without the addition of estrogen and were randomly assigned.**

- Data should be collected and processed randomly or appropriately blocked. **Data from different conditions were collected and processed simultaneously under identical circumstances.**

#### Blinding

- Allocation concealment: The investigator should be unaware of the group to which the next animal taken from a cage will be allocated. **The investigator was unaware of the group allocation.**

- Blinded conduct of the experiment: Animal caretakers and investigators conducting the experiments should be blinded to the allocation sequence. **The animal caretakers were blinded to the allocation sequence of the experiments.**

- Blinded assessment of outcome: Investigators assessing, measuring, or quantifying experimental outcomes should be blinded to the intervention. **The investigator assessed the outcomes partly blinded. Blinding was done for the immunocytochemical analyses that were manually counted.**

#### Sample-size Estimation

- An appropriate sample size should be computed when the study is being designed, and the statistical method of computation reported. **Post-hoc power calculations have been conducted, showing a power of 82%. This indicates that our study had a high probability of detecting real differences between the groups, suggesting that our sample sizes and experimental design were adequate.**

- Statistical methods that take into account multiple evaluations of the data should be used when an interim evaluation is carried out. **We did not perform interim evaluations of the data. All statistical analyses were conducted only after data collection was completed to ensure unbiased results.**

#### Data Handling

- Rules for stopping data collection should be defined in advance. **We did not stop data collection; all data were collected as planned. During the analysis phase,**

**we assessed whether the neuronal networks met the predefined inclusion criteria.**

- Criteria for inclusion and exclusion of data should be established prospectively. **Only neuronal networks demonstrating good quality, such as sufficient cell density for proper neuron-electrode coupling and an even distribution of cells, were included. For inclusion in the analysis, neuronal networks needed to display a mean firing rate (MFR) greater than 0.1 spikes per second and at least 1 network burst per minute.**

- How outliers will be defined and handled should be decided when the experiment is being designed, and any data removed before analysis should be reported. **Outliers were identified using the Robust Regression and Outlier Removal (ROUT) method with Q = 1%, resulting in the removal of data points in the lower and upper 1% of the normal distribution.**

- The primary endpoint should be prospectively selected. If multiple endpoints are to be assessed, then appropriate statistical corrections should be applied. **We investigated the time evolution of different parameters, but did not perform direct comparisons between these parameters. Each parameter was analyzed independently, so no statistical corrections for multiple comparisons were required.**

- Investigators should report on data missing because of attrition or exclusion. **We initially started with a total of 120 neuronal networks on MEAs across all experimental conditions. Specifically, we used 36 male control, 36 female control, 24 male estrogen-treated, and 24 female estrogen-treated neuronal networks. After excluding some networks based on pre-established inclusion criteria, the final number of neuronal networks included in the analysis was 22 for MC, 29 for FC, 13 for ME, and 19 for FE. We also started with 120 neuronal networks on coverslips. We used 32 male control and 32 female control networks, with 8 assigned to each time point (normoxia, 8h, 16h, 24h, and 36h of hypoxia). We used 16 networks each for male and female estrogen-treated groups, with 4 networks assigned to each time point. Networks were excluded based on inclusion criteria. Numbers of included coverslips are reported in supplementary Table 8.**

### Pseudo-replication

**We considered pseudo-replication by averaging outcomes per parameter for each experimental unit and analyzing temporal evolution by assessing the best model to describe the data. Since multiple measurements from the same unit were not treated as independent data points, pseudo-replication was avoided in our study design and analysis.**

#### Repetition of Experiments

**For the control groups, data were collected from 4 separate experiments. For the estrogen groups, data were collected from 2 separate experiments. All experiments were conducted under identical conditions in a computer-controlled climate chamber to ensure consistency. Each experiment involved separate cortex isolations.**
